# Identification and characterization of novel plasma proteins in drug resistant HIV/AIDS patients by SWATH-MS

**DOI:** 10.1101/2021.03.04.433855

**Authors:** Sushanta Kumar Barik, Srikanth Prasad Tripathy, Deepa Bisht, Praveen Singh, Rahul Chakraborty, Monu Kumar Chadar, Shripad A Patil, Tej Pal Singh, Rekha Tandon, Srikanta Jena, Keshar Kunja Mohanty

**Author notes:** Correspondence address, Dr Keshar Kunja Mohanty, Scientist-F, National JALMA Institute for Leprosy and Other Mycobacterial Diseases, Taj-Ganj, Agra, Uttar-Pradesh, India, Email ID.

## Abstract

**Background and Objectives:** Acquired immunodeficiency syndrome is one of the most important diseases caused by human immunodeficiency virus. Understanding its molecular pathogenesis is essential to manage the disease at the population level. In this study, a quantitative analysis of plasma proteins was carried out in drug resistant and drug respondent patients using the SWATH-MS.

**Methods:** Sequential window acquisition of all theoretical mass spectra (SWATH-MS) is a prime technique to seek the key plasma proteins involved in virus replication and drug metabolism during therapy.

**Results:** In total, 204 proteins were identified and quantified, 57 proteins were differentially expressed, 25 proteins were down regulated and 32 proteins were upregulated in drug resistant patients. Six proteins such as complement C4-A, immunoglobulin heavy variable 1-2, carboxylic ester hydrolase, fibulin-1, immunoglobulin lambda constant 7, secreted phosphoprotein 24 were statistically expressed in drug resistant patients compared to the drug respondent patients. Gene ontology study and protein-protein interaction networks were established in six statistically significant differentially expressed proteins of the drug resistant patients.

**Interpretation and Conclusion:** Our findings high lights the novel proteins that were differentially expressed in drug resistant patients. A label-free quantitative proteomics method for depleted human plasma samples by SWATH-MS that can be useful in plasma proteomics research in any biological system.

## Introduction

Plasma proteins play very important role in a variety of diseases. Protease inhibitors were known to bind efficiently to plasma proteins and 90% bound to Alfa-1 acid glycoprotein. Efavirenz, a non-nucleoside reverse transcriptase inhibitor (NNRTI) binds to plasma proteins and 99% bound to albumin. The nucleoside reverse transcriptase inhibitors (NRTIs) were not highly bound to plasma proteins. Thus, protein binding influences the antiviral activity of the drugs ^1^. The analysis of HIV-1 infected human plasma is an attractive medium with which differentially expressed proteins could be identified in certain disease states. Proteomic techniques are widely used globally to identify differentially expressed plasma proteins in response to HIV-1 infection. Sequential window acquisition of all theoretical mass spectra (SWATH-MS) is a novel technique conceptualized with the protein library and the individual protein was identified and quantified for biomarker discovery ^2,3^ .The study elaborates with the plasma proteomics of HIV-1 patients treated with first-line ART over 6 years. The main search was that the various proteins and their fold changes associated with multiple therapies in drug-resistant and drug-respondent patients.

## Materials and Methods

### Materials

Ammonium formate, formic acid, dithiothreitol (DTT) and iodoacetamide (IAA) were procured from Sigma, USA. Modified trypsin (sequencing grade) was procured from Promega, USA. Polysulfoethyl SCX cartridge (5 micron, 300 Å bead) with cartridge holder was procured from Sciex, USA. Acetonitrile, Liquid phase-chromatography-mass spectroscopy grade water, was procured from Biosolve (Lorraine, France). Analytical grade chemicals were used in this study.

### Study design and Study participants

The study was focused on six first-line ART such as ZLE, ZLN, TLE, TLN, SLE and SLN patients who were enrolled in the anti-retroviral therapy centre, Sarojini Naidu Medical College, Agra, India from December 2009 to November 2016 as per the treatment guidelines directed by NACO, Govt. of India and ethical guidelines by ICMR, Govt. of India ^4-6^. The age ranges of six patients were from 32 years to 43 years. A patient information leaflet was used for data collection ^7^.

Ten ml of blood samples was collected from each patient whose CD4^+^ count was <350 cells/µl. Then plasma was extracted after centrifuging at 2000g for 10 min. Then plasma was stored at -80°C for viral load and genotyping at NIRT, ICMR, Chennai, Tamil-Nadu, India.

The viral load was estimated in all six samples. Among them, 3 patients had a higher viral load >1000 copies/ml (drug resistant) and 3 patients had a lower viral load <1000 copies/ml to target not detected level (drug respondent).Three drug-resistant patients (>1000 copies/ml) were considered for genotyping.

### Ethics committee and informed consent

The name of the Ethics Committee is Institute Human Ethics Committee of National JALMA Institute for Leprosy and Other Mycobacterial Diseases, (Indian Council of Medical Research), Dr. M. Miyazaki Marg, Tajganj, Agra-282004. The registration number of institute is Institute Human Ethics Committee was ECR/257/Inst/ UP/ 2013. The ethics committee meeting was held on 27.5.2016 at National JALMA Institute for Leprosy and Other Mycobacterial Diseases, Agra for the project entitled “Characterization of drug resistant HIV-1 mutants of Agra region, India by genomic and proteomic approaches”. The registration number of the Institute Human Ethics Committee was ECR/257/Inst/ UP/ 2013as per the guidelines directed by the Indian Council of Medical Research, Govt. of India^4.^ Renewal of registration of Institute Human Ethics Committee of National JALMA Institute for leprosy and Other Mycobacterial Diseases was done by Central Drugs Standard Control Organization (CDSCO) with registration no. ECR/257/Inst/UP/2013/RR-20 dated 19^th^ June 2020 and provisional registration with department of Health Research, Ministry of Health and Family Welfare, Govt. of India, File number-EC/NEW/INST/2020/792 dated 27^th^ July 2020.The written informed consent was obtained from the study participant.

### Genotyping

The genotyping was performed with samples with a viral load ≥1000 copies/ml at the National Institute for Research in Tuberculosis, Chennai, by using the WHO dried blood spot protocol 2010 ^8^. The details of the PCR primers and reaction conditions are reported earlier ^9^.

### Protein characterization by SWATH-MS

#### Sample preparation for proteomics analysis

The 100µl plasma was inactivated by heating at 56°C for 30 min ^10^ .60µl plasma was used for albumin and globulin depletion using an Aurum serum mini kit (BioRad, USA). The six depleted plasma samples were used for SWATH-MS analysis. Protein estimation was done on the depleted plasma protein samples using the Bradford assay (Sigma-Aldrich, USA). In the SWATH-MS analysis following steps like reduction, alkylation and trypsin digestion, Library generation for SWATH analysis, data searches and peak extraction have been conducted by the published protocol of Ghose et al, 2019, Basak et al, 2015 ^11,12^. The detail of SWTH-MS protocol is given in the supplementary file-S6.

### Statistical analysis

The processed .mrkvw files from PeakView were then loaded onto MarkerView (version 1.2.1, AB Sciex) and total area sum normalisation was performed, protein peak areas were then exported in excel where statistical analysis (Student’s t-test) was performed. Proteins with a 1.5-fold increase or decrease (p□value < 0.05) were taken as differential proteins.

## Results

Blood samples from three drug resistant patients having a viral load of 9983 copies/ml,12344 copies/ml, 36177 copies/ml respectively and three drug respondent patients having a viral load of <40 copies/ml, target not detected level, 47 copies/ml respectively were analysed for SWATH MS study. The NRTIs and NNRTIs associated mutations M41L, T215Y, M184V, T215F, and Y188L, G190A were identified in three drug-resistant patients.

### Protein folds change analysis of drug-resistant and drug-respondent patients

Retention time (RT) calibration of peptides and peak area extraction was performed in the SWATH micro app of PeakView and data normalization was done in MarkerView. Peak areas were then exported to excel and then statistical analysis was performed. The data were normalized using total area sum normalization. The normalized fold change over resistant to respondent was calculated and >1.5 value was considered as significantly upregulated proteins and <0.67 value for significantly downregulated proteins. Finally, a student t-test was done to compare between drug resistant and drug respondent patients for each detected protein. P<0.05 was considered significant.

Thirty-two proteins were identified as upregulated and twenty-five proteins were identified as down regulated in drug resistant patients.

The heat map of the differentially expressed proteins was designed based on the Z score ^13.^The expression profile of each differentially expressed proteins are presented in the heat map in the figure-1.

**Figure-1:**
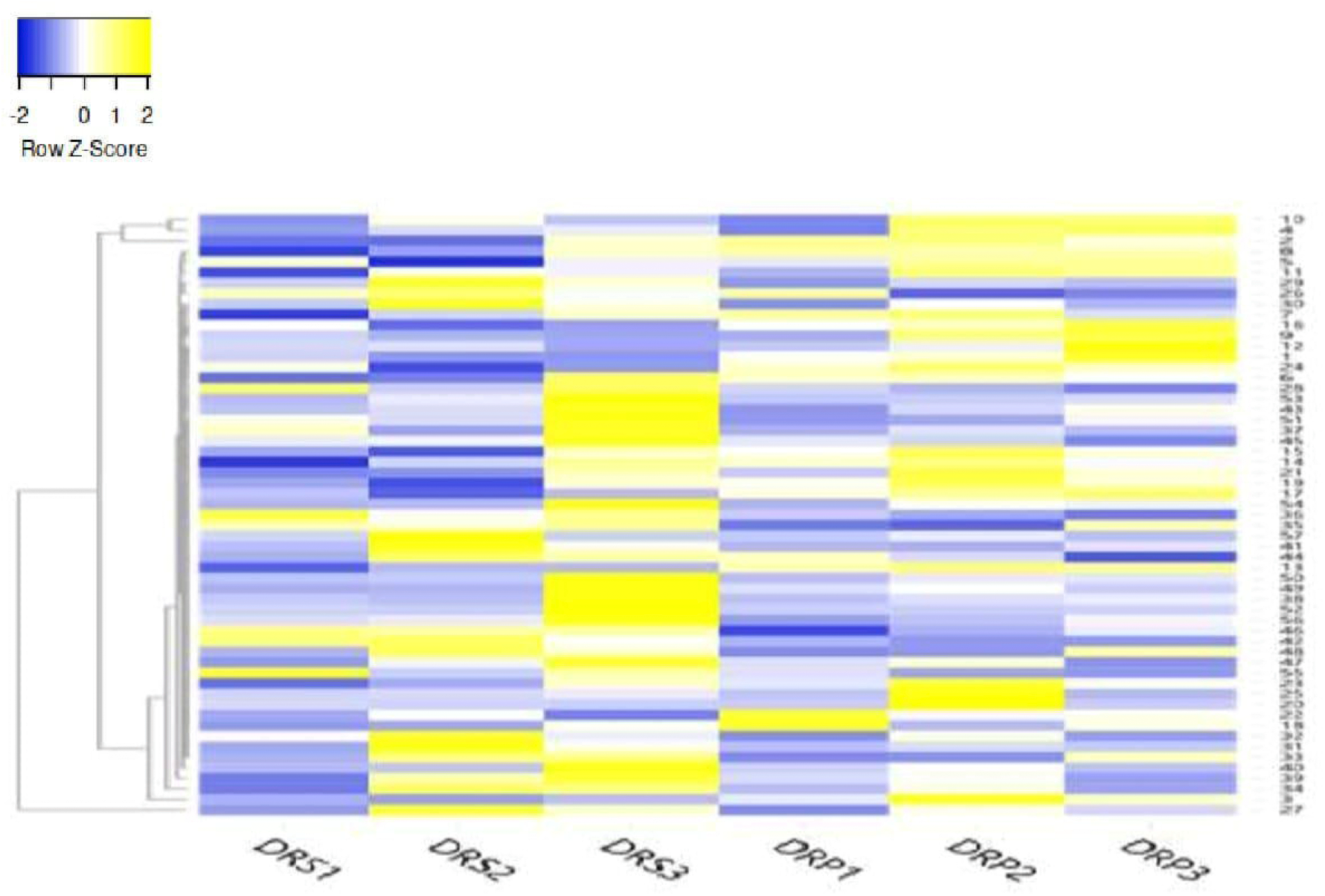
Heat map of the differentially expressed down regulated and upregulated proteins DRS represents Drug resistant and DRP represents Drug respondent. The expression of each protein is based on Z score range from -2 (blue colour) to 2 (Yellow colour). The serial number of the down regulated and upregulated proteins in the heat map are represented in the Gene symbol: l.C4A 2. IGHM 3. APOA4 4.HBB, 5. SERPINGl 6. PZP 7. IGHG2 8.CD5L 9.HPR 10. HBAl 11. JCHAIN 12. APOLl 13. IGHVl-2 14. IGHV3-9 15. IGHV5-5116. C4B 17. BCHE 18. LPA 19. IGLV6-57 20. SERPINAl 21. IGKV2-24 22. IGLV3-10 23. FAT-1 24.BIN3 25. PROC 26. FNl 27. IGHAl 28. GSN 29. AFM 30. TTR 31. IGLV3-21 32. KRTl 33. IGLV3-19 34. IGKV3-15 35. C4BPB 36. FBLNl 37. IGKVl-5 38. F5 39. IGKV3-ll 40. IGLV7-46 41. VNNl 42. IGLC7 43. IGHA2 44. IGHV3-13 45. IGHV1ORl5-l 46. SPP2 47. CDKL3 48. IGLVl0-54 49. CDC42BPA 50. SP5 51. BEX5 52. PEPD 53. IGHV6-l 54. IGHVl-45 55. CDH5 56. 3 57. SERPINA 10 The heat map of differentially expressed proteins were created by the online tool (http://www.heatmapper.ca/expression).

### Bioinformatic analyses

All 57-protein gene symbol and ID were retrieved through Uniprot (www.uniprot.org). The details of the molecular functions of the 57 differentially expressed proteins in the drug resistant patients was analysed using PANTHER-gene analyst ^14^. The molecular functions of the differentially expressed proteins are presented in the figure-2.

**Figure-2:**
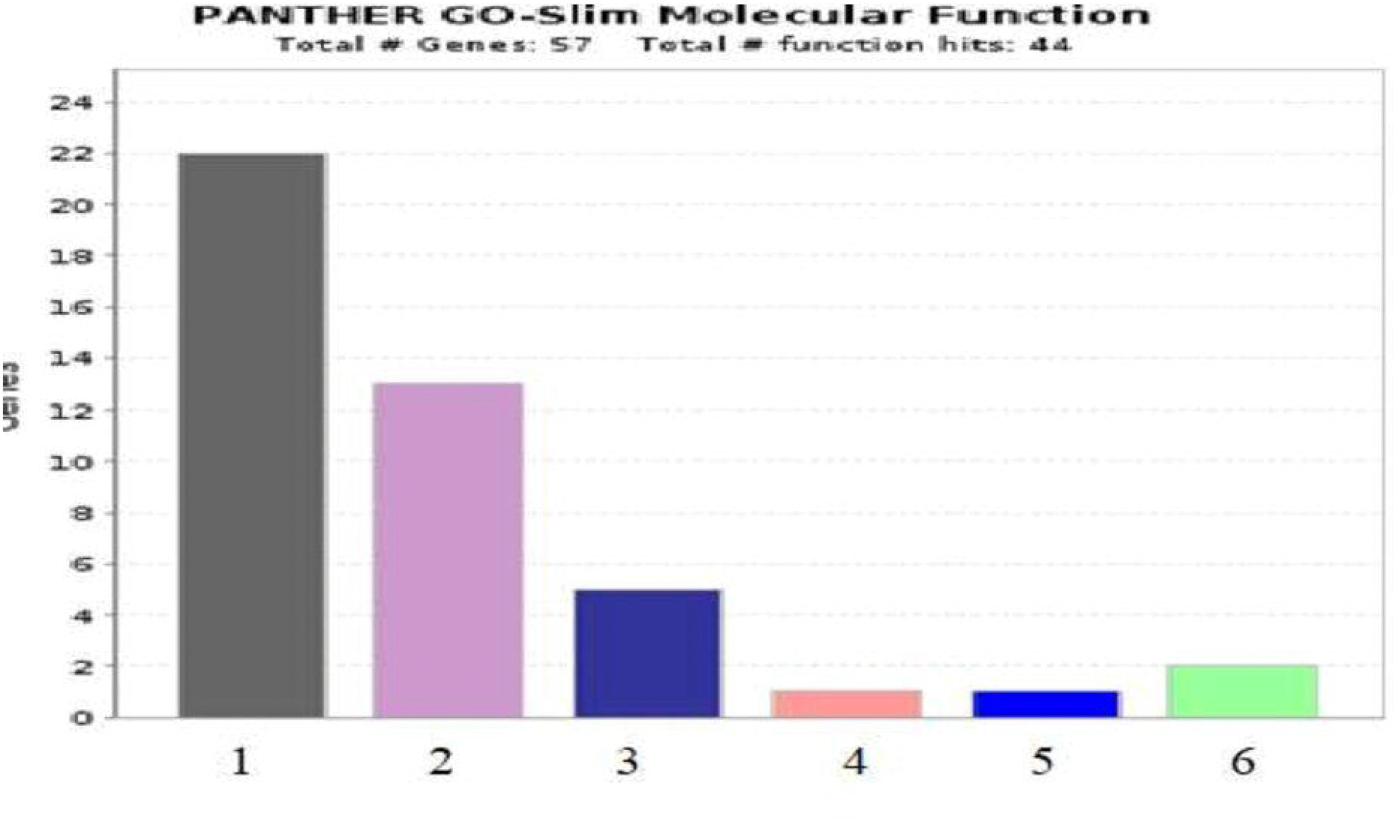
Molecular functions of the differential expressed proteins-**PANTHER**-gene analyst. The gene list for a category of the molecular function of the differential expressed proteins are presented in the bar diagram. The number of genes involved in the different functional role is presented in serial number in the bar diagram 1. Binding (GO:0005488), 2. Catalytic activity (GO:0003824), 3. Molecular function regulator (GO:0098772), 4. Molecular transducer activity (GO:0060089), 5. Transcription regulator activity (GO:0140110), 6. Transporter activity (GO:0005215). The molecular function of the differential expressed proteins was created by the online tool **PANTHER-**gene analyst (http://www.pantherdb.org).

Analysing the panther pathway component list and panther pathway suggested that few plasma proteins were involved in important pathways like the FAS signalling pathway, blood coagulation pathway, cadherin signaling pathway, B-cell activation, Wnt signaling pathway, plasminogen activating cascade, integrin signaling pathway, nicotinic acetyl choline receptor signaling pathway, muscarinic acetyl choline receptor 1 and 3 signaling pathway, muscarinic acetyl choline receptor 2 and 4 signaling pathway, Alzheimer disease-presenilin pathway and CCKR signalling map of HIV-1 patients. The major identified 22 plasma proteins in HIV-1 patients were involved in the binding activity in the panther molecular function list. The detail of the panther pathway analysis is given in the supplementary table S1.

The Reactome pathway database (https://reactome.org/) was used ^15^. The 25 most relevant pathways were inter connected with these 57 differentially expressed proteins in the plasma of drug resistant patients. The details of the Reactome pathways are given in a supplementary table S2. Gene enrichment analysis of 57 differential expressed proteins encoded by genes of the drug resistant patients was performed by Fun Rich (http://www.funrich.org/) ^16^. In the Biological pathway, a set of genes (29.41%) was involved in homeostasis, signalling events mediated by VEGFR1 and VEGFR2, VEGF and VEGFR signalling network, Sphingosine 1-phosphate (S1P) pathway, integrin family cell surface interactions and FAS (CD95) signalling pathway. In biological processes, a set of genes was involved in (23.33%) cell growth and maintenance (20.0%) and immune response (16.66%). Regarding the molecular function, a set of genes were involved in protease inhibitor activity (13.33%) and transporter activity (10.0%). The site of expression of a set of genes (93.54%) was in the plasma of drug resistant patients. The transcription factor HNF4A regulates the expression of a set of genes (47.82%) in the plasma of drug resistant patients. The protein domain of a set of genes (72.41%) acts as signal peptide. In the clinical phenotype, a set of genes (90.90%) was found to be involved in inherited

diseases like autosomal dominant in the drug resistant patients. The differentially expressed proteins are presented in the volcano plot in the Fig.3. The volcano plot was made using Graph Pad prism8,0 (Graph pad prism, USA) ^17^.

**Figure-3:**
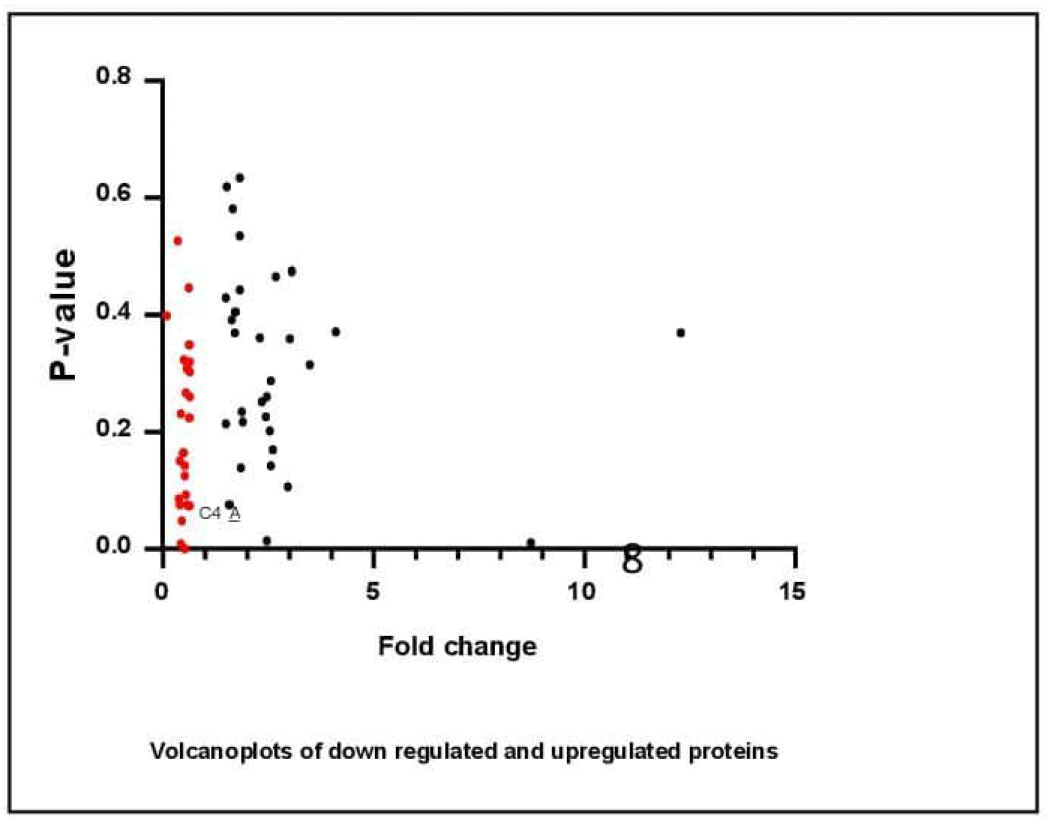
Volcano plots of down -regulated and up-regulated proteins The red colour dots represent the down-regulated proteins and black colour dots represent the up-regulated proteins (Graph pad Prism 8, USA). The volcano plot was created by Graph Pad Prism 8, USA(https://www.graphpad.com/scientific-software/prism/).

All 57 differentially expressed proteins encoded by genes of the drug resistant patients was analysed using the Kyoto Encyclopaedia of Genes and Genomes (KEGG) (https://www.genome.jp/) ^18^. In the KEGG pathway, these 57 proteins were involved in 28 pathways, interacted with two networks and expressed in 26 diseases. The details of the KEGG analysis are given in the supplementary table S3.

### Protein-protein interaction network

Protein-protein interactions among all the differentially expressed plasma proteins were analysed using STRING v.11 (STRING) ^19^. Protein-protein interactions were investigated and link was shown with the functional partners (Fig.4).

**Figure-4:**
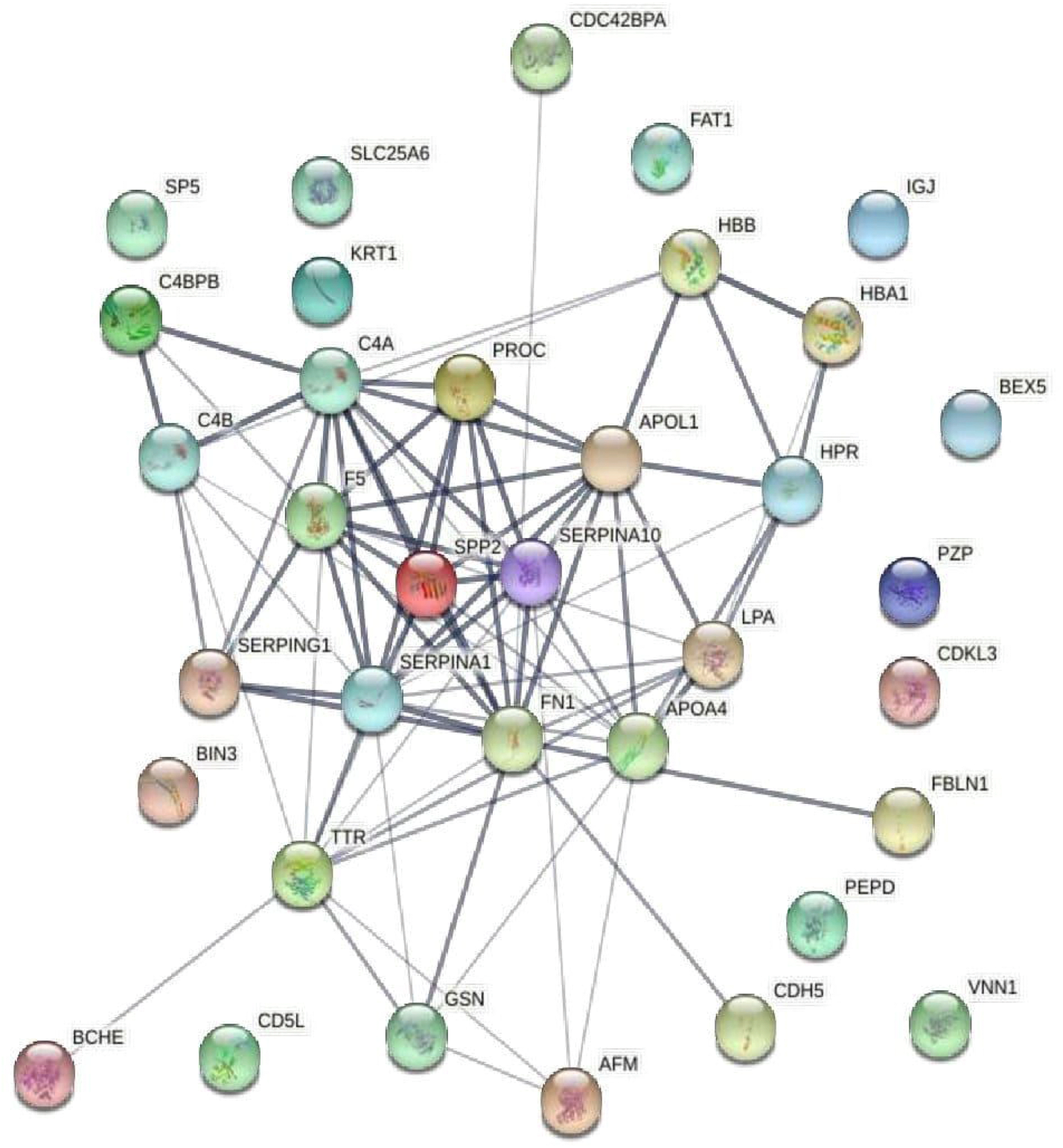
Protein-protein interaction network The details description of the protein-protein interaction network nodes represents proteins: Splice isoforms or post translational modifications are coUapsed i.e. Each node represents alJ the proteins produced by a single protein-coding gene locus. The edges represent protein-protein associations: Network status :number of nodes: 35, number of edges:80, average node degree: 4.57, average local clustering coefficient:0.453, expected number of edges: 7, PPI enrichment p-value:< l.0e-16. The protein-protein interaction network was created by using the online tool STRING Version 11.0 (https://string-db.org).

### Gene ontology of statistically differentially expressed proteins

The statistically significantly differentially expressed three proteins, such as complement C4-A (fold change=0.45, P=0.05), immunoglobulin heavy variable 1-2 (fold change=0.53, P=0.005), carboxylic ester hydrolase (fold change=0.43, P=0.01) were down regulated and the three proteins fibulin-1(fold change=1.82, P=0.03), immunoglobulin lambda constant 7 (fold change=8.7, P=0.01), secreted phosphoprotein 24 (fold change=2.4, P=0.01) were upregulated in the drug-resistant patients. Gene ontology such as molecular functions, biological processes and cellular components of the statistically significant differentially expressed proteins were analysed by STRINGv.11 (STRING) and described in supplementary table S4. Gene set enrichment analysis of six statistically differentially expressed proteins were analysed using Fun Rich 3.1.3. In Biological pathway, a set of statistically differentially expressed genes (33.33%) was involved in the initial triggering of complement, synthesis, secretion, and diacylation of ghrelin, complement cascade, innate immune system, epithelial-to-mesenchymal transition, diabetes pathways, and immune system. In biological processes, a set of genes (25%) was involved in immune response, cell growth /maintenance, protein metabolism, energy pathways, and metabolism. All proteins were extracellular. Molecular function analysis of a set of genes (25%) indicates the complement activity, protease inhibitor activity, extracellular matrix structural constituent, hydrolase activity etc. All sets of genes encoded proteins in the plasma. LHX3 and HNF4A transcription factors were regulating the set of genes (100%) in the plasma of drug resistant patients. Statistically significant differentially expressed genes were analysed through the KEGG pathway (www.kegg.jp).

### Pathway list by KEGG

□ hsa04610: Complement and coagulation cascades - Homo sapiens (human)
□ hsa05322: Systemic lupus erythematosus - Homo sapiens (human)
□ hsa05133: Pertussis - Homo sapiens (human)
□ hsa05150: Staphylococcus aureus infection - Homo sapiens (human)

### Diseases

□ H00080: Systemic lupus erythematosus
□ H00102: Classic complement pathway component defects
□ H01649: Schizophrenia

### Drug and protein interaction study

The six-drug respondent and drug resistant patients were in first-line ART such as ZLE, ZLN, SLN, TLE, and TLN during their course of treatment in this period. The drug interactions with these proteins were analysed (www.dgidb.org) ^20^. A total of 44 proteins were identified those which are interacting with zidovudine, lamivudine, stavudine, nevirapine, and efavirenz drugs. Tenofovir interaction with any protein was not observed in this drug interaction database. The detailed analysis of protein-drug interaction file is given in a supplementary table S5.

## Discussion

The differential expression and molecular function of the Serotransferrin and Apolipoprotein A1 was identified in HIV-1 patients treated with first line ART ^9^. Now, the fundamental role of these six statistically significant differentially expressed proteins such as complement C4-A, immunoglobulin heavy variable 1-2, carboxylic ester hydrolase, fibulin-1, immunoglobulin lambda constant 7, secreted phosphoprotein 24 and their association with several human diseases were discussed.

The complement system is activated in the human plasma after HIV-1 infection. Once the complement is activated after infection, the outcome would be virolysis. The intricacies between complement and antibodies were reviewed during the course of HIV infection ^21^. C4 is an important component in the complement system that plays important role in defence mechanism. C4 deficiency is associated with several bacterial and viral diseases.C4a is an allotype of C4gene. C4a protein is an anaphylatoxin family of proteins produced by the activation of complement. We found, C4a protein is downregulated in drug-resistant HIV-1 patients. It can be considered that the down regulation of C4a can increase the viral copy number in drug-resistant HIV-1 patients. Down regulation of the C4a protein may not activate the complement system and virolysis may not happen after activation of the complement system. Even the patients are in first-line anti-retroviral therapy, the drugs would not work to kill the virus due to NRTIs and NNRTIs associated mutations in the RT gene of HIV-1. Therefore, this may be hypothesized that the RT gene mutation and downregulation of C4a have the impacts on increased viral copy number in drug-resistant HIV-1 patients.

Human antibodies play an important in the control of HIV-1 infection. The strategy for boosting the early HIV-1 specific IgG response should include in early antiretroviral therapy and trials of therapeutic vaccines in HIV patients ^22^. The immunoglobulin heavy chain variable region (IGHV) is responsible for antigen binding and comprised of HIV-1 individuals who developed neutralizing antibodies ^23^. The variable region of immunoglobulin heavy variable 1-2 was participated in antigen recognition. Immunoglobulins are antibodies secreted by B-lymphocytes. This variable region of immunoglobulin eliminates the bound antigen and an effector molecule in humoral immunity ^24^. Thus, the down regulation of the immunoglobulin heavy variable 1-2 may affect the binding activity and elimination of the HIV particles in these drug -resistant HIV-1 patients.

Carboxylic ester hydrolase plays an important role in the hydrolysis of some drugs into inactive metabolites in human plasma. The study of the role of esterase would be helpful in drug development and clinical pharmacotherapy ^25^. The oral antiretrovirals first hydrolysed and then absorbed into the gastrointestinal tract. The implications of safe drug therapy are so important in HIV-1 patients. The down regulation of the carboxylic ester hydrolase may not hydrolyse the drugs in the case of these drug -resistant HIV-1 patients.

Fibulin-1 act as a tumour suppressor gene and angiogenesis inhibitor in bladder cancer. Fibulin-1 was epigenetically down regulated in bladder cancer. Fibulin-1 down regulation was associated with the non-muscle invasive bladder cancer grade and recurrence ^26^. However, the fibulin-1 is up-regulated in the plasma of dug resistant HIV-1 patients by the SWATH-MS data analysis. These three-drug resistant HIV-1 patients were failure with first line ART over 1 to 6 years. The expression of fibulin-1 is probably the novel finding in drug resistant HIV-1 patients. The implication of this finding is yet to be explored.

The immunoglobulin lambda chain gene was encoded by two separate germ line genes such as specificity region gene and common region gene, expressed a single continuous polypeptide chain. The main role of Immunoglobulin λ chain is antigen binding in human infection and shown the biased properties. A significant bias toward use of the λ light chain in the anti-HIV Env response was observed in individuals with acute HIV infection, those with chronic HIV infection, HIV-negative vaccines and HIV clades (clade B, clade G, and CRF02_AG).The biased λ chain is associated with enhanced binding of anti-HIV Env glycoprotein antibodies in HIV patients and an effector molecule in humoral response ^27^. The protein immunoglobulin lambda constant 7 is upregulated in drug resistant HIV-1 patients and this upregulated expression in plasma may not compete in the enhanced binding of anti-HIV Env glycoprotein and may not act as an effector molecule in humoral response in drug resistant patients.

Secreted phosphoprotein is a member of cystatin superfamily encoded by SPP2 gene. Secreted phosphoprotein 24 is a bone matrix protein secreted from the liver. The secreted phosphoprotein 24 regulates bone metabolism ^28^. Bone morphogenetic proteins bind to and affect the activity of bone morphogenetic proteins. SPP24 protein regulates the formation and administration of the activity of the TGF-β during bone growth and development ^29^. In our study, we observed the secreted phosphoprotein 24 is upregulated in drug-resistant patients that may affect the bone metabolism in drug resistant HIV-1 patients.

These major findings of the six novel plasma proteins in drug resistant patients will add value in the basic proteomic study by SWATH-MS. In future, studying the mechanism of these novel proteins by in vitro and in vivo experiments would throw some light in the life cycle of HIV-1 and drug metabolism in human plasma.

## Conclusion

The identification and quantification of the significantly differentially expressed proteins by SWATH -MS is a novel approach. Possibly for the first time we are reporting these proteins identified in drug resistant and drug respondent HIV-1 patients who are taking first line ART over 6 years. Attempts have been made to specify the role of these proteins through gene ontology study. This study would be helpful for further implementation in the search for biomarker discovery in HIV-1 patients.

The whole proteomics data generated by SWATH-MS were submitted into the PRIDE. The project title and accession number are given below:

Project Name: Characterization of plasma proteins of HIV/AIDS patients under first-line anti-retroviral therapy, Project accession: PXD016843.

## Acknowledgements

Indian Council of Medical Research, Govt. of India is acknowledged.

## Financial support and sponsorship

The project “Characterization of drug resistant HIV-1 mutants of Agra region, India by genomic and proteomic approaches” funded for senior research fellowship of Mr Sushanta Kumar Barik (File No. 80/990/2015-ECD-I). Mr M. M. Alam, JALMA, ICMR and Mr Rakesh Kumar Mishra, ART centre, S.N Medical College is acknowledged for providing technical help in the collection of blood samples.

## Conflict of interest

All authors declare no conflict of interest.

## Notes

### Competing Interest Statement

The authors have declared no competing interest.

