## supplemental table1 for "Identification and characterization of novel plasma proteins in drug resistant HIV/AIDS patients by SWATH-MS"

Supplementary Table S1

**Table S1: Panther gene list, Panther pathway component list and Panther Pathway List**

**Panther gene list:**

| Sl No. | **Gene ID** | **Mapped ID** | **Gene name**  **Gene symbol** | **Panther family/subfamily** | **Panther protein class** | **Species** |
| --- | --- | --- | --- | --- | --- | --- |
| 1 | HUMAN\|HGNC=19953\|  UniProtKB=Q8IWX5 | SPP2 | Sphingosine-1-phosphate phosphatase 2  SGPP2 ortholog | SPHINGOSINE-1- PHOSPHATE PHOSPHATASE 2 (PTHR14969:SF14) | Phosphatase | Homo  sapiens |
| 2 | HUMAN\|HGNC=15483\|  UniProtKB=Q8IVW4 | CDKL3 | Cyclin-dependent kinase-like3 CDKL3 ortholog | CYCLIN-DEPENDENT KINASE-LIKE-3 (PTHR24056:SF177) | Non-receptor serine/threonine protein kinase H | Homo  sapiens |
| 3 | HUMAN\|HGNC=5930\|  UniProtKB=A0A075B6I9 | IGLV7-46 | Immunoglobulin lambda variable 7-46 IGLV7-46 ortholog | IMMUNOGLOBULIN LAMBDA VARIABLE7-46 (PTHR23267:SF120) | Immunoglobulin | Homo  sapiens |
| 4 | HUMAN\|HGNC=1764\|  UniProtKB=P33151 | CDH5 | Cadherin-5 CDH5 ortholog | CADHERIN-5 (PTHR24027:SF89) | **--------------** | Homo  sapiens |
| 5 | HUMAN\|HGNC=3542\|  UniProtKB=P12259 | F5 | Coagulation factor V  F5 ortholog | COAGULATION FACTORV (PTHR24543:SF302) | oxidoreductase | Homo  sapiens |
| 6 | HUMAN\|HGNC=5741\|  UniProtKB=P01602 | IGKV1-5 | Immunoglobulin kappa variable 1- 5  IGKV1-5 ortholog | IMMUNOGLOBULIN KAPPA VARIABLE 1-5  (PTHR23267:SF373) | Immunoglobulin | Homo  sapiens |
| 7 | HUMAN\|HGNC=5581\|  UniProtKB=P01766 | IGHV3-13 | Immunoglobulin heavy variable 3- 13  IGHV3-13 ortholog | IMMUNOGLOBULIN HEAVY VARIABLE 3-13 (PTHR23266:SF316) | Immunoglobulin | Homo  sapiens |
| 8 | HUMAN\|HGNC=5816\|  UniProtKB=P01624 | IGKV3-15 | Immunoglobulin kappa variable 3- 15  IGKV3-15 ortholog | IMMUNOGLOBULIN KAPPA VARIABLE 3-15-RELATED (PTHR23267:SF331) | Immunoglobulin | Homo  sapiens |
| 9 | HUMAN\|HGNC=15996\|  UniProtKB=Q9UK55 | SERPINA10 | Protein Zdependent protease inhibitor SERPINA10 ortholog | PROTEINZ-DEPENDENT PROTEASE INHIBITOR (PTHR11461:SF191) | Protease inhibitor | Homo  sapiens |
| 10 | HUMAN\|HGNC=8840\|UniProtKB=P12955 | PEPD | Xaa-Pro dipeptidase PEPD ortholog | XAA-PRO DIPEPTIDASE (PTHR43226:SF1) | Metalloprotease | Homo  sapiens |
| 11 | HUMAN\|HGNC=12705\|UniProtKB=O95497 | VNN1 | Pantetheinase VNN1 ortholog | PANTETHEINASE (PTHR10609:SF16) | Hydrolase | Homo  sapiens |
| 12 | HUMAN\|HGNC=1737\|UniProtKB=Q5VT25 | CDC42BPA | Serine/threonineprotein kinase MRCK alpha CDC42BPA ortholog | SERINE/THREONINE-PROTEIN KINASE MRCK ALPHA (PTHR22988:SF31) | Non-receptor serine/threonine protein kinase | Homo  sapiens |
| 13 | HUMAN\|HGNC=11256\|UniProtKB=Q13103 | SPP2 | Secreted phosphoprotein 24  SPP2 ortholog | SECRETED PHOSPHOPROTEIN24 (PTHR15444:SF4) | **-------------------** | Homo  sapiens |
| 14 | HUMAN\|HGNC=5550\|UniProtKB=P23083 | IGHV1-2 | Immunoglobulin heavy variable 1- 2 IGHV1-2 ortholog | IMMUNOGLOBULIN HEAVY VARIABLE 1-2 (PTHR23266:SF272) | Immunoglobulin | Homo  sapiens |
| 15 | HUMAN\|HGNC=5884\|UniProtKB=A0A075B6I4 | IGLV10-54 | Immunoglobulin lambda variable 10-54 IGLV10-54 ortholog | IMMUNOGLOBULIN LAMBDA VARIABLE 10-54 (PTHR23267:SF225) | Immunoglobulin | Homo  sapiens |
| 16 | HUMAN\|HGNC=4620\|UniProtKB=P06396 | GSN | Gelsolin GSN ortholog | GELSOLIN-RELATED (PTHR11977:SF29) | Non-motor actin binding protein | Homo  sapiens |
| 17 | HUMAN\|HGNC=5526\|  UniProtKB=P01859 | IGHG2 | Immunoglobulin heavy constant gamma 2 IGHG2 ortholog | IMMUNOGLOBULIN HEAVY CONSTANT GAMMA 1-RELATED (PTHR23266:SF83) | Immunoglobulin | Homo  sapiens |
| 18 | HUMAN\|HGNC=1690\|  UniProtKB=O43866 | CD5L | CD5 antigen-like CD5L ortholog | CD5 ANTIGEN-LIKE (PTHR48071:SF8) - | **------------** | Homo  sapiens |
| 19 | HUMAN\|HGNC=5478\|  UniProtKB=P01876 | IGHA1 | Immunoglobulin heavy constant alpha 1 IGHA1 ortholog | IMMUNOGLOBULIN HEAVY CONSTANT ALPHA 1-RELATED (PTHR23266:SF68) | Immunoglobulin | Homo  sapiens |
| 20 | HUMAN\|HGNC=618\|  UniProtKB=O14791 | APOL1 | Apolipoprotein L1 APOL1 ortholog | APOLIPOPROTEIN L1 (PTHR14096:SF55) | Apolipoprotein | Homo  sapiens |
| 21 | HUMAN\|HGNC=4823\|  UniProtKB=P69905 | HBA1 | Hemoglobin subunit alpha HBA2 ortholog | HEMOGLOBIN SUBUNIT ALPHA (PTHR11442:SF48) - | **----------------** | Homo  sapiens |
| 22 | HUMAN\|HGNC=27990\|UniProtKB=Q5H9J7 | BEX5 | Protein BEX5 BEX5 ortholog | PROTEIN BEX5 (PTHR19430:SF0) | **---------------** | Homo  sapiens |
| 23 | HUMAN\|HGNC=1054\|UniProtKB=Q9NQY0 | BIN3 | Bridging integrator 3 BIN3 ortholog | BRIDGING INTEGRATOR 3 (PTHR47174:SF3) | **-------------** | Homo  sapiens |
| 24 | HUMAN\|Gene=KV311_HUMAN\|UniProtKB=P04433 I | IGKV3-11 | Immunoglobulin kappa variable 3- 11 IGKV3-11 ortholog | IMMUNOGLOBULIN KAPPA VARIABLE 3- 11-RELATED (PTHR23267:SF344) | Immunoglobulin | Homo  sapiens |
| 25 | HUMAN\|HGNC=1328\|UniProtKB=P20851 | C4BPB | C4b-binding protein beta chain C4BPB ortholog | C4B-BINDING PROTEIN BETA CHAIN-RELATED (PTHR45656:SF9) | **--------------** | Homo  sapiens |
| 26 | HUMAN\|HGNC=5861\|UniProtKB=A0M8Q6 | IGLC7 | Immunoglobulin lambda constant 7 IGLC7 ortholog | IMMUNOGLOBULIN LAMBDA CONSTANT 7 (PTHR23266:SF296) |  | Homo  sapiens |
| **27** | HUMAN\|HGNC=5479\|UniProtKB=P01877 | IGHA2 | Immunoglobulin heavy constant alpha 2 IGHA2 ortholog | IMMUNOGLOBULIN HEAVY CONSTANT ALPHA 1-RELATED (PTHR23266:SF68) | Immunoglobulin | Homo  sapiens |
| 28 | HUMAN\|HGNC=5563\|UniProtKB=A0A075B7D0 | IGHV1OR15- 1 | Immunoglobulin heavy variable 1/OR15-1 (nonfunctional) (Fragment) IGHV1OR15-1 ortholog | IMMUNOGLOBULIN HEAVY VARIABLE 1-2  (PTHR23266:SF272) | Immunoglobulin | Homo  sapiens |
| 29 | HUMAN\|HGNC=5541\|UniProtKB=P01871 | IGHM | Immunoglobulin heavy constant mu IGHM ortholog | IMMUNOGLOBULIN HEAVY CONSTANT MU (PTHR23266:SF120) | Immunoglobulin | Homo  sapiens |
| 30 | HUMAN\|HGNC=8941\|UniProtKB=P01009 | SERPINA1 | Alpha-1- antitrypsin SERPINA1 ortholog | ALPHA-1- ANTITRYPSIN (PTHR11461:SF165) | Protease inhibitor | Homo  sapiens |
| 31 | HUMAN\|HGNC=5713\|UniProtKB=P01591 | JCHAIN | Immunoglobulin J chain JCHAIN ortholog | IMMUNOGLOBULIN J CHAIN (PTHR10070:SF2) | Immunoglobulin | Homo  sapiens |
| 32 | HUMAN\|HGNC=9750\|UniProtKB=P20742 | PZP | Pregnancy zone protein PZP ortholog | PREGNANCY ZONE PROTEIN (PTHR11412:SF92) | Protease inhibitor | Homo  sapiens |
| 33 | HUMAN\|HGNC=602\|  UniProtKB=P06727 | APOA4 | Apolipoprotein AIV APOA4 ortholog | APOLIPOPROTEIN AIV (PTHR18976:SF1) | **------------** | Homo  sapiens |
| 34 | HUMAN\|HGNC=1228\|UniProtKB=P05155 | SERPING1 | Plasma protease C1 inhibitor SERPING1 ortholog | PLASMA PROTEASE C1 INHIBITOR (PTHR11461:SF159) | Protease inhibitor | Homo  sapiens |
| 35 | HUMAN\|HGNC=5659\|UniProtKB=A0A0C4DH38 I | IGHV5-51 | Immunoglobulin heavy variable 5- 51 IGHV5-51 ortholog | IMMUNOGLOBULIN HEAVY VARIABLE 5-10-1-RELATED (PTHR23266:SF322) | Immunoglobulin | Homo  Sapiens |
| 36 | HUMAN\|HGNC=12405\|  UniProtKB=P02766 | TTR | Transthyretin TTR ortholog | TRANSTHYRETIN (PTHR10395:SF12) | Hydrolase | Homo  Sapiens |
| 37 | HUMAN\|HGNC=3595\|UniProtKB=Q14517 | FAT1 | Protocadherin Fat 1 FAT1 ortholog | PROTOCADHERIN FAT 1 (PTHR24026:SF42) | Cadherin | Homo  Sapiens |
| 38 | HUMAN\|HGNC=5927\|  UniProtKB=P01721 | IGLV6-57 | Immunoglobulin lambda variable 6-57 IGLV6-57 ortholog | IMMUNOGLOBULIN LAMBDA VARIABLE 6-57 (PTHR23267:SF108) | Immunoglobulin | Homo  Sapiens |
| 39 | HUMAN\|HGNC=5553\|UniProtKB=A0A0A0MS14 I | IGHV1-45 | Immunoglobulin heavy variable 1- 45  IGHV1-45 ortholog | IMMUNOGLOBULIN HEAVY VARIABLE 1- 45 (PTHR23266:SF285) | Immunoglobulin | Homo  sapiens |
| 40 | HUMAN\|HGNC=5897\|UniProtKB=A0A075B6K4 | IGLV3-10 | Immunoglobulin lambda variable 3-10 IGLV3-10 ortholog | IMMUNOGLOBULIN LAMBDA VARIABLE 3-10 (PTHR23267:SF322) | Immunoglobulin | Homo  sapiens |
| 41 | HUMAN\|HGNC=1323\|UniProtKB=P0C0L4 | C4A | Complement C4-A  C4A ortholog | COMPLEMENT C4-ARELATED (PTHR11412:SF86) | Protease inhibitor | Homo  sapiens |
| 42 | HUMAN\|Gene=HV309_HUMAN\|UniProtKB=P01782 | IGHV3-9 | Immunoglobulin heavy variable 3- 9 IGHV3-9 ortholog | IMMUNOGLOBULIN HEAVY VARIABLE 3-9 (PTHR23266:SF324) | Immunoglobulin | Homo  sapiens |
| 43 | HUMAN\|HGNC=14529\|UniProtKB=Q6BEB4 | SP5 | Transcription factor Sp5  SP5 ortholog | TRANSCRIPTION FACTOR SP5 (PTHR23235:SF29) | C2H2 zinc finger transcription factor | Homo  sapiens |
| 44 | HUMAN\|HGNC=6667\|  UniProtKB=P08519 | LPA | Apolipoprotein(a) LPA ortholog | APOLIPOPROTEIN(A)-RELATED (PTHR24261:SF2) | Serine protease | Homo  sapiens |
| 45 | HUMAN\|HGNC=316\|  UniProtKB=P43652 | AFM | Afamin AFM ortholog | AFAMIN (PTHR11385:SF14) | Transfer/carrier protein | Homo  sapiens |
| 46 | HUMAN\|HGNC=983\|  UniProtKB=P06276 | BCHE | Cholinesterase  BCHE ortholog | CHOLINESTERASE (PTHR43918:SF5) | **------------** | Homo  sapiens |
| 47 | HUMAN\|HGNC=5156\|  UniProtKB=P00739 | HPR | Haptoglobinrelated protein HPR ortholog | HAPTOGLOBINRELATED (PTHR24255:SF27) | Serine protease | Homo  sapiens |
| 48 | HUMAN\|HGNC=9451\|  UniProtKB=P04070 | PROC | Vitamin -K dependent protein C PROC ortholog | VITAMIN K-DEPENDENT PROTEIN C (PTHR24278:SF0) | Serine  protease | Homo  sapiens |
| 49 | HUMAN\|HGNC=10802\|  UniProtKB=P11686 | SP5 | Pulmonary surfactantassociated protein C SFTPC ortholog | PULMONARY SURFACTANTASSOCIATED PROTEIN C (PTHR10800:SF4) | Surfactant | Homo  sapiens |
| 50 | HUMAN\|HGNC=5662\|  UniProtKB=A0A0B4J1U7 | IGHV6-1 | Immunoglobulin heavy variable 6- 1 IGHV6-1 ortholog | IMMUNOGLOBULIN HEAVY VARIABLE 4- 34-RELATED (PTHR23266:SF238) | Immunoglobulin | Homo  sapiens |
| 51 | HUMAN\|HGNC=4827\|UniProtKB=P68871 | HBB | Hemoglobin subunit  beta HBB ortholog | HEMOGLOBIN SUBUNIT BETA (PTHR11442:SF42) | **---------** | Homo  sapiens |
| 52 | HUMAN\|HGNC=5781\|  UniProtKB=A0A0C4DH68 | IGKV2-24 | Immunoglobulin kappa variable 2- 24 IGKV2-24 ortholog | IMMUNOGLOBULIN KAPPA VARIABLE 2- 24 (PTHR23267:SF379) | Immunoglobulin | Homo  sapiens |
| 53 | HUMAN\|HGNC=6412\|  UniProtKB=P04264 | KRT1 | Keratin, type II cytoskeletal 1  KRT1 ortholog | KERATIN, TYPE II CYTOSKELETAL 1 (PTHR45616:SF33) |  | Homo  sapiens |
| 54 | HUMAN\|HGNC=5905\|UniProtKB=P80748 | IGLV3-21 | Immunoglobulin lambda variable 3-21 IGLV3-21 ortholo | IMMUNOGLOBULIN LAMBDA VARIABLE 3-21 (PTHR23267:SF378) |  | Homo  sapiens |
| 55 | HUMAN\|HGNC=3778\|UniProtKB=P02751 | FN1 | Fibronectin FN1 ortholog | FIBRONECTIN (PTHR19143:SF267) | Intercellular signal molecule | Homo  sapiens |
| 56 | HUMAN\|HGNC=3600\|UniProtKB=P23142 | FBLN1 | Fibulin-1 FBLN1 ortholog | FIBULIN-1 (PTHR24050:SF23) - |  | Homo  sapiens |
| 57 | HUMAN\|HGNC=5903\|UniProtKB=P01714 | IGLV3-19 | Immunoglobulin lambda variable 3-19 IGLV3-19 | IMMUNOGLOBULIN LAMBDA VARIABLE 3-19 (PTHR23267:SF364) | Immunoglobulin | Homo  sapiens |

**Panther pathway list**

| **Pathway**  **Accession** | **Mapped ID** | **Pathway**  **Name** | **Components** | **Subfamilies** | **Associated sequence** |
| --- | --- | --- | --- | --- | --- |
| P00011 | P00408 P00426 P00415 P00435 P00425 P00423 P00421 P00432 | Blood coagulation | 59 | 47 | 205 |
| P00020 | P00611 | Fas signalling pathway | 31 | 46 | 243 |
| P06959 | G06981 P07171 G07274 | CCKR signaling map | 290 | 171 | 171 |
| P00004 | P00118 P00117 P00139 P00168 P00165 | Alzheimer disease presenillin pathway | 70 | 208 | 1037 |
| P00057 | P01440 | Wnt signaling pathway | 49 | 494 | 2296 |
| P00012 | P00471 | Cadherin signnaling pathway | 16 | 237 | 1054 |
| P00050 | P01246 P01255 | Plasminogen activating cascade | 22 | 18 | 82 |
| P00042 | P01067 | Muscarinic acetylcholine receptor 1 and 3 signalling pathway | 12 | 96 | 507 |
| P00010 | P00389 | B cell activation | 37 | 80 | 427 |
| P00043 | P01080 | Muscarinic acetylcholine receptor1 and3 signalling pathway | 11 | 112 | 523 |
| P00044 | P01093 | Nicotinic acetylcholine receptor signalling pathway | 13 | 245 | 959 |
| P00034 | P00939 | Integrin signalling pathway | 46 | 261 | 1295 |

**Panther Pathway Component List**

| Component mapped IDs  Accession | Component Name  Type | Upstream | Down stream | Pathway |
| --- | --- | --- | --- | --- |
| [P00611](http://www.pantherdb.org/pathway/pathCatDetail.do?clsAccession=P00611)  HUMAN\|HGNC=4620\|UniProtKB=P06396 | Gesolin  Protein |  | Protein [Caspase-3](http://www.pantherdb.org/pathway/pathCatDetail.do?clsAccession=P00599) | FAS signaling  pathway |
| [P00421](http://www.pantherdb.org/pathway/pathCatDetail.do?clsAccession=P00421)  HUMAN\|HGNC=8941\|UniProtKB=P01009 | [Alpha1-antitrypsin](http://www.pantherdb.org/pathway/pathCatDetail.do?clsAccession=P00421)  Protein |  | [Thrombin](http://www.pantherdb.org/pathway/pathCatDetail.do?clsAccession=P00414) | Blood coagulation |
| [P00471](http://www.pantherdb.org/pathway/pathCatDetail.do?clsAccession=P00471)  HUMAN\|HGNC=3595\|UniProtKB=Q14517  HUMAN\|HGNC=1764\|UniProtKB=P33151 | [Cadherin](http://www.pantherdb.org/pathway/pathCatDetail.do?clsAccession=P00471)  Protein |  |  | Cadherin  signaling  pathway |
| [P00432](http://www.pantherdb.org/pathway/pathCatDetail.do?clsAccession=P00432)  HUMAN\|HGNC=3542\|UniProtKB=P12259 | FV  Protein | [Thrombin](http://www.pantherdb.org/pathway/pathCatDetail.do?clsAccession=P00414) |  | Blood coagulation |
| [P00435](http://www.pantherdb.org/pathway/pathCatDetail.do?clsAccession=P00435)  HUMAN\|HGNC=3542\|UniProtKB=P12259 | FVa  Protein |  |  | Blood coagulation |
| [P00425](http://www.pantherdb.org/pathway/pathCatDetail.do?clsAccession=P00425)  HUMAN\|HGNC=15996\|UniProtKB=Q9UK55 | [ZPI](http://www.pantherdb.org/pathway/pathCatDetail.do?clsAccession=P00425)  Protein |  | [Factor Xia](http://www.pantherdb.org/pathway/pathCatDetail.do?clsAccession=P00434) | Blood coagulation |
| [P00389](http://www.pantherdb.org/pathway/pathCatDetail.do?clsAccession=P00389)  HUMAN\|HGNC=5541\|UniProtKB=P01871 | [mIgM](http://www.pantherdb.org/pathway/pathCatDetail.do?clsAccession=P00389)  Protein | [Lyn](http://www.pantherdb.org/pathway/pathCatDetail.do?clsAccession=P00374) |  | B cell activation |
| [P01093](http://www.pantherdb.org/pathway/pathCatDetail.do?clsAccession=P01093)  HUMAN\|HGNC=983\|UniProtKB=P06276 | [AChE](http://www.pantherdb.org/pathway/pathCatDetail.do?clsAccession=P01093)  Protein |  |  | Nicotinic  acetylcholine  receptor sign |
| [P01246](http://www.pantherdb.org/pathway/pathCatDetail.do?clsAccession=P01246)  HUMAN\|HGNC=6667\|UniProtKB=P08519 | Plasmin  Protein |  | [Tissue type plasminogen activator](http://www.pantherdb.org/pathway/pathCatDetail.do?clsAccession=P01257) [pro-matrix metalloprotease 13](http://www.pantherdb.org/pathway/pathCatDetail.do?clsAccession=P01254)  Matrix metalloprotease 1  Fibrin pro-Matrix metalloprotease 1  pro-Matrix metalloprotease 3  pro-matrix metalloprotease 9  Matrix metalloprotease 13  Matrix metalloprotease 9  Matrix metalloprotease 3 | Plasminogen  activating  cascade |
| P01067  HUMAN\|HGNC=983\|UniProtKB=P06276 | AChE  Protein |  |  | Muscarinic  acetylcholine  receptor 1 |
| P00426  HUMAN\|HGNC=9451\|UniProtKB=P04070 | PC  Protein |  |  | Blood coagulation |
| P00939  HUMAN\|HGNC=3778\|UniProtKB=P02751 | Fibronectin  Protein |  |  | Integrin signalling  pathway |
| P01255  HUMAN\|HGNC=6667\|UniProtKB=P08519 | Plasminogen  Protein | Tissue type plasminogen activator - |  | Plasminogen  activating  cascade |
| P01080  HUMAN\|HGNC=983\|UniProtKB=P06276 | AChE  Protein |  |  | Muscarinic  acetylcholine  receptor 2 |
| P01440  HUMAN\|HGNC=3595\|UniProtKB=Q14517 HUMAN\|HGNC=1764\|UniProtKB=P33151 | Cadherin  Protein |  |  | Wnt signaling  pathway |
| P00423  HUMAN\|HGNC=9451\|UniProtKB=P04070 | APC  Protein |  |  | Blood  coagulation |

The panther class, panther Pathway component list and panther pathway list were analyzed by the online tool

PANTHER-gene list analysis ( http://www.pantherdb.org) database (<https://reactome.org/>) was used.
