## Supplemental table2 for "Identification and characterization of novel plasma proteins in drug resistant HIV/AIDS patients by SWATH-MS"

**Supplementary table S2**

Table S2: Reactome pathways

The following table shows the 25 most relevant pathways sorted by p-value.

| **Scavenging of heme from plasma** 19 / 106 | 0.007 | 1.11e-16 | 6.66e-15 | 6 / 12 | 9.45e-04 |
| --- | --- | --- | --- | --- | --- |
| **Regulation of Complement cascade** 17 / 139 | 0.01 | 1.11e-16 | 6.66e-15 | 18 / 42 | 0.003 |
| **Complement cascade** 17 / 156 | 0.011 | 1.11e-16 | 6.66e-15 | 27 / 71 | 0.006 |
| **Initial triggering of complement** 15 / 120 | 0.008 | 1.11e-16 | 6.66e-15 | 6 / 21 | 0.002 |
| **Binding and Uptake of Ligands by** 19 / 167 | 0.012 | 1.11e-16 | 6.66e-15 | 6 / 33 | 0.003 |
| **Cell surface interactions at the** 18 / 257 | 0.018 | 3.33e-16 | 1.67e-14 | 4 / 64 | 0.005 |
| **Classical antibody-mediated** 13 / 97 | 0.007 | 1.78e-15 | 7.64e-14 | 2 / 2 | 1.57e-04 |
| **FCGR activation** 13 / 103 | 0.007 | 3.77e-15 | 1.40e-13 | 6 / 6 | 4.72e-04 |
| **Creation of C4 and C2 activators** 13 / 111 | 0.008 | 9.66e-15 | 3.19e-13 | 2 / 8 | 6.30e-04 |
| **Role of phospholipids in** 13 / 129 | 0.009 | 6.39e-14 | 1.92e-12 | 5 / 12 | 9.45e-04 |
| **CD22 mediated BCR regulation** 11 / 72 | 0.005 | 7.41e-14 | 2.00e-12 | 3 / 4 | 3.15e-04 |
| **Role of LAT2/NTAL/LAB on calcium** 12 / 107 | 0.007 | 1.85e-13 | 4.47e-12 | 2 / 7 | 5.51e-04 |
| **FCGR3A-mediated IL10 synthesis** 13 / 141 | 0.01 | 1.95e-13 | 4.47e-12 | 9 / 20 | 0.002 |
| **Hemostasis** 24 / 824 | 0.057 | 3.37e-13 | 7.07e-12 | 41 / 328 | 0.026 |
| **FCGR3A-mediated phagocytosis** 13 / 157 | 0.011 | 7.42e-13 | 1.26e-11 | 12 / 27 | 0.002 |
| **Parasite infection** 13 / 157 | 0.011 | 7.42e-13 | 1.26e-11 | 12 / 27 | 0.002 |
| **Leishmania phagocytosis** 13 / 157 | 0.011 | 7.42e-13 | 1.26e-11 | 12 / 27 | 0.002 |
| **Regulation of actin dynamics for** 13 / 158 | 0.011 | 8.03e-13 | 1.28e-11 | 6 / 24 | 0.002 |
| **FCERI mediated MAPK activation** 12 / 124 | 0.009 | 1.02e-12 | 1.53e-11 | 2 / 20 | 0.002 |
| **FCERI mediated Ca+2 mobilization** 12 / 129 | 0.009 | 1.60e-12 | 2.41e-11 | 2 / 11 | 8.66e-04 |
| **Antigen activates B Cell Receptor**  **(BCR) leading to generation of** 11 / 103  **second messengers** | 0.007 | 3.39e-12 | 4.74e-11 | 10 / 25 | 0.002 |
| **Fcgamma receptor (FCGR)** 13 / 193 | 0.013 | 9.51e-12 | 1.24e-10 | 17 / 42 | 0.003 |
| **Vesicle-mediated transport** 22 / 824 | 0.057 | 2.31e-11 | 3.01e-10 | 26 / 251 | 0.02 |
| **FCERI mediated NF-kB activation** 12 / 175 | 0.012 | 5.25e-11 | 6.31e-10 | 1 / 19 | 0.001 |
| **Innate Immune System** 25 / 1,329 | 0.092 | 1.14e-09 | 1.37e-08 | 63 / 697 | 0.055 |

| **Pathway name** | **Entities** | | | | **Reactions** | |
| --- | --- | --- | --- | --- | --- | --- |
|  | **found** | **ratio** | **p-value** | **FDR*** | **found** | **ratio** |

* False discovery rate

The reactome pathways were analysed by the online toll (<https://reactome.org> )
