## supplemental table3 for "Identification and characterization of novel plasma proteins in drug resistant HIV/AIDS patients by SWATH-MS"

Supplementary table S3

**Table S3: KEGG pathway list**

```
hsa04610 Complement and coagulation cascades - Homo sapiens (human) (7)
hsa05143 African trypanosomiasis - Homo sapiens (human) (4)
hsa05133 Pertussis - Homo sapiens (human) (4)

hsa05322 Systemic lupus erythematosus - Homo sapiens (human) (2)
hsa05150 Staphylococcus aureus infection - Homo sapiens (human) (2)
hsa05144 Malaria - Homo sapiens (human) (2)
hsa04979 Cholesterol metabolism - Homo sapiens (human) (2)

hsa04810 Regulation of actin cytoskeleton - Homo sapiens (human) (2)
hsa04977 Vitamin digestion and absorption - Homo sapiens (human) (1)
hsa04512 ECM-receptor interaction - Homo sapiens (human) (1)
 hsa04666 Fc gamma R-mediated phagocytosis - Homo sapiens (human) (1)
 hsa05146 Amoebiasis - Homo sapiens (human) (1)
hsa04918 Thyroid hormone synthesis - Homo sapiens (human) (1)

hsa04933 AGE-RAGE signaling pathway in diabetic complications - Homo sapiens (human) (1)
hsa04670 Leukocyte transendothelial migration - Homo sapiens (human) (1)
hsa00770 Pantothenate and CoA biosynthesis - Homo sapiens (human) (1)
hsa04510 Focal adhesion - Homo sapiens (human) (1)
hsa05418 Fluid shear stress and atherosclerosis - Homo sapiens (human) (1)
hsa05222 Small cell lung cancer - Homo sapiens (human) (1)
hsa05200 Pathways in cancer - Homo sapiens (human) (1)

hsa05165 Human papillomavirus infection - Homo sapiens (human) (1)
hsa04151 PI3K-Akt signaling pathway - Homo sapiens (human) (1)
hsa05135 Yersinia infection - Homo sapiens (human) (1)
hsa04514 Cell adhesion molecules (CAMs) - Homo sapiens (human) (1)
 hsa04975 Fat digestion and absorption - Homo sapiens (human) (1)
hsa05100 Bacterial invasion of epithelial cells - Homo sapiens (human) (1)
 hsa05205 Proteoglycans in cancer - Homo sapiens (human) (1)
hsa05203 Viral carcinogenesis - Homo sapiens (human) (1)
```

**KEGG network**

[06135](https://www.genome.jp/kegg-bin/show_network?id=06135&amp;queryfile=searh_network.84249.args&amp;align=1) Cytoskeletal regulation (viruses) (1)

[06167](https://www.genome.jp/kegg-bin/show_network?id=06167&amp;queryfile=searh_network.84249.args&amp;align=1) Human cytomegalovirus (HCMV) (1)

**Diseases**

```
H00102 Classic complement pathway component defects (2) [path:hsa04610] H00106 Complement regulatory protein defects (2)
H00845 Familial amyloidosis (1)

H00798 Familial carpal tunnel syndrome (1)
H01006Hereditary angioedema (1) [path:hsa04610] H01433 Budd-Chiari syndrome (1)
H01723 Deep vein thrombosis (1)
H00228 Thalassemia (2)
H00229 Sickle cell anemia (1)

H00220 Factor V deficiency (1) [path:hsa04610]
H00223 Inherited thrombophilia (2) [path:hsa04610]
H01103 Alpha-1-antitrypsin deficiency (1) [path:hsa04610]
H01714 Chronic obstructive pulmonary disease (COPD) (1)
H00626 Focal segmental glomerulosclerosis (1)
H01260 Glomerulopathy with fibronectin deposits (1) [path:hsa04510 hsa04512 hsa04810]
H01119 Prolidase deficiency (1)
H00459 Synpolydactyly (1)

H02185 Spondylometaphyseal dysplasia (1)

H00691 Bullous congenital ichthyosiform erythroderma (BCIE) (1)
H00707 Ichthyosis hystrix, Curth-Macklin type (1)
H02265 Annular epidermolytic ichthyosis (1)
     H00717 Striate palmoplantar                         keratoderma (1)
       H00722Epidermolytic palmoplantar keratoderma (1)
       H01649 Schizophrenia (1)
        H02322 Amyloidosis, Finnish type (1)
```

The KEGG pathway was analysed by the online tool (<https://www.genome.jp/kegg/pathway.html>).
