## supplemental table4 for "Identification and characterization of novel plasma proteins in drug resistant HIV/AIDS patients by SWATH-MS"

Supplementary table S4.

**Table S4 : Details of gene ontology of six significant statistically differential expressed proteins**

**Functional enrichments in your set of proteins**

**Biological Process (GO)**

*GO-term	description	count in gene set	false discovery rate*

[GO:0052547](http://amigo.geneontology.org/amigo/term/GO:0052547)	regulation of peptidase activity	3 of 420	0.0114

[GO:0072376](http://amigo.geneontology.org/amigo/term/GO:0072376)	protein activation cascade	2 of 74	0.0129

[GO:0032269](http://amigo.geneontology.org/amigo/term/GO:0032269) negative regulation of cellular

protein metabolic process	3 of 1014	0.0390

[GO:0010951](http://amigo.geneontology.org/amigo/term/GO:0010951) negative regulation of

endopeptidase activity	2 of 242	0.0442

**Molecular Function (GO)**

*GO-term	description	count in gene set	false discovery rate*

[GO:0061134](http://amigo.geneontology.org/amigo/term/GO:0061134)	peptidase regulator activity	3 of 211	0.00025

[GO:0004866](http://amigo.geneontology.org/amigo/term/GO:0004866)	endopeptidase inhibitor activity	2 of 169	0.0108

**Cellular Component (GO)**

*GO-term	description	count in gene set	false discovery rate*

[GO:0005788](http://amigo.geneontology.org/amigo/term/GO:0005788)	endoplasmic reticulum lumen	3 of 299	0.00094

[GO:0005576](http://amigo.geneontology.org/amigo/term/GO:0005576)	extracellular region	4 of 2505	0.0088

**Reactome Pathways**

*Pathway	description	count in gene set	false discovery rate*

[HSA-8957275](https://reactome.org/content/detail/R-HSA-8957275) Post-translational protein

Phosphorylation	2 of 106	0.0049

[HSA-381426](https://reactome.org/content/detail/R-HSA-381426) Regulation of Insulin-like

Growth Factor (IGF) transport

and uptake by Insulin-like Growth

Factor Binding Proteins (IGFBPs )	2 of 123	0.0049

[HSA-392499](https://reactome.org/content/detail/R-HSA-392499)	Metabolism of proteins	3 of 1948	0.0330

**INTERPRO Protein Domains and Features**

*Domain	description	count in gene set	false discovery rate*

[IPR000020](https://www.ebi.ac.uk/interpro/entry/IPR000020)	Anaphylatoxin/fibulin	2 of 6	2.54e-05

**SMART Protein Domains**

*Domain	description	count in gene set	false discovery rate*

[SM00104](http://smart.embl-heidelberg.de/smart/do_annotation.pl?DOMAIN=SM00104) Anaphylatoxin homologous

Domain	2 of 6	6.14e-06

hsa05322	Systemic lupus erythematosus	2 of 94	0.0454

Gene ontology of six statistically differentially expressed proteins were analysed by the online tool STRING V.11(<https://string-db.org/>).
