## supplemental table5 for "Identification and characterization of novel plasma proteins in drug resistant HIV/AIDS patients by SWATH-MS"

**Supplementary table S5**

**Table S5: Protein-drug interaction**

| search_term | match_term | match_type | gene | drug | Interaction types | sources | pmids |
| --- | --- | --- | --- | --- | --- | --- | --- |
| ZIDOVUDINE | ZIDOVUDINE | Definite | TERT | ZIDOVUDINE | inhibitor | Drug Bank | 23303810 |
| ZIDOVUDINE | ZIDOVUDINE | Definite | ABCB1 | ZIDOVUDINE |  | NCI | 15456083 |
| ZIDOVUDINE | ZIDOVUDINE | Definite | GZMA | ZIDOVUDINE |  | NCI | 12820184 |
| ZIDOVUDINE | ZIDOVUDINE | Definite | PRL | ZIDOVUDINE |  | NCI | 3877233\|9500147 |
| ZIDOVUDINE | ZIDOVUDINE | Definite | CXCL10 | ZIDOVUDINE |  | NCI | 11141242 |
| ZIDOVUDINE | ZIDOVUDINE | Definite | LDLR | ZIDOVUDINE |  | NCI | 8211145 |
| ZIDOVUDINE | ZIDOVUDINE | Definite | ABCC4 | ZIDOVUDINE |  | Pharm GKB\|NCI | 12105214 |
| ZIDOVUDINE | ZIDOVUDINE | Definite | CD4 | ZIDOVUDINE |  | NCI | 1998495 |
| ZIDOVUDINE | ZIDOVUDINE | Definite | KRT18 | ZIDOVUDINE |  | NCI | 17713161 |
| ZIDOVUDINE | ZIDOVUDINE | Definite | IL10 | ZIDOVUDINE |  | NCI | 10433557 |
| ZIDOVUDINE | ZIDOVUDINE | Definite | UGT2B7 | ZIDOVUDINE |  | Pharm GKB |  |
| ZIDOVUDINE | ZIDOVUDINE | Definite | ENOSF1 | ZIDOVUDINE |  | NCI | 12238529\|15980332 |
| ZIDOVUDINE | ZIDOVUDINE | Definite | TERC | ZIDOVUDINE |  | NCI | 11036953 |
| ZIDOVUDINE | ZIDOVUDINE | Definite | PTGS1 | ZIDOVUDINE |  | NCI | 18049160 |
| ZIDOVUDINE | ZIDOVUDINE | Definite | IL18 | ZIDOVUDINE |  | NCI | 12559970 |
| LAMIVUDINE | LAMIVUDINE | Definite | ENOSF1 | LAMIVUDINE |  | NCI | 10864683 |
| LAMIVUDINE | LAMIVUDINE | Definite | HPRT1 | LAMIVUDINE |  | NCI | 17358033 |
| LAMIVUDINE | LAMIVUDINE | Definite | ABCC1 | LAMIVUDINE |  | NCI | 12218384 |
| LAMIVUDINE | LAMIVUDINE | Definite | APOA1 | LAMIVUDINE |  | NCI | 17538878 |
| LAMIVUDINE | LAMIVUDINE | Definite | CYP2B6 | LAMIVUDINE |  | Pharm GKB |  |
| LAMIVUDINE | LAMIVUDINE | Definite | MMP1 | LAMIVUDINE |  | NCI | 15309715 |
| LAMIVUDINE | LAMIVUDINE | Definite | LEP | LAMIVUDINE |  | NCI | 16091072 |
| LAMIVUDINE | LAMIVUDINE | Definite | ABCC4 | LAMIVUDINE |  | Pharm GKB |  |
| LAMIVUDINE | LAMIVUDINE | Definite | PTH | LAMIVUDINE |  | NCI | 16171525 |
| STAVUDINE | STAVUDINE | Definite | BRD4 | STAVUDINE | inhibitor | GuideTo PharmacologyInteractions | |
| STAVUDINE | STAVUDINE | Definite | CSF2 | STAVUDINE |  | NCI | 2538549 |
| STAVUDINE | STAVUDINE | Definite | CYP2B6 | STAVUDINE |  | Pharm GKB |  |
| STAVUDINE | STAVUDINE | Definite | DTYMK | STAVUDINE |  | NCI | 15134538 |
| STAVUDINE | STAVUDINE | Definite | CXCL10 | STAVUDINE |  | NCI | 11141242 |
| STAVUDINE | STAVUDINE | Definite | TNFRSF1B | STAVUDINE |  | NCI | 9430255 |
| EFAVIRENZ | EFAVIRENZ | Definite | NR1I2 | EFAVIRENZ |  | Pharm GKB |  |
| EFAVIRENZ | EFAVIRENZ | Definite | NR1I3 | EFAVIRENZ |  | Pharm GKB |  |
| EFAVIRENZ | EFAVIRENZ | Definite | CYP2B6 | EFAVIRENZ |  | NCI\|FDA | 17041008 |
| EFAVIRENZ | EFAVIRENZ | Definite | CYP3A4 | EFAVIRENZ |  | NCI | 16176117 |
| NEVIRAPINE | NEVIRAPINE | Definite | CYP2B6 | NEVIRAPINE |  | NCI | 17041008 |
| NEVIRAPINE | NEVIRAPINE | Definite | ADIPOQ | NEVIRAPINE |  | NCI | 15305892 |
| NEVIRAPINE | NEVIRAPINE | Definite | CYP2A7P1 | NEVIRAPINE |  | PharmGKB |  |
| NEVIRAPINE | NEVIRAPINE | Definite | PSORS1C2 | NEVIRAPINE |  | PharmGKB |  |
| NEVIRAPINE | NEVIRAPINE | Definite | CCHCR1 | NEVIRAPINE |  | PharmGKB |  |
| NEVIRAPINE | NEVIRAPINE | Definite | APOB | NEVIRAPINE |  | NCI | 12869587 |
| NEVIRAPINE | NEVIRAPINE | Definite | MBLAC2 | NEVIRAPINE |  | PharmGKB |  |
| NEVIRAPINE | NEVIRAPINE | Definite | ABCC10 | NEVIRAPINE |  | PharmGKB |  |
| NEVIRAPINE | NEVIRAPINE | Definite | CYP2C19 | NEVIRAPINE |  | PharmGKB |  |
| NEVIRAPINE | NEVIRAPINE | Definite | CYP3A4 | NEVIRAPINE |  | PharmGKB |  |

The protein -drug interaction study was analysed by the online tool ( [www.dgidb.org](http://www.dgidb.org)).
