## supplemental table6 for "Identification and characterization of novel plasma proteins in drug resistant HIV/AIDS patients by SWATH-MS"

**Supplementary file S6**

**SWATH-MS protocol for plasma proteins**

**Reduction, alkylation and trypsin digestion**

For SWATH run, 10µg of proteins from each sample were reduced with 25mM DTT for 25 min at 56°C, followed by alkylation using 55mM IAA at room temperature for 15-20 min in the dark, and trypsin digestion for 18 h at 37°C. Tryptic peptides were vacuum-dried in a vacuum concentrator.

**LC-MS/MS data acquisition**

**Library generation for SWATH analysis**

For generation of the spectral ion library in data-dependent acquisition mode, 300 µg of protein obtained by pooling of plasma from the six drug resistant and drug respondent patients were digested using trypsin as described earlier (Ghose et al., 2019)^11^ and were fractionated into 8 fractions by cation exchange (SCX) chromatography using a SCX cartridge (5 micron, 300 Å bead from AB Sciex, USA) and a step-gradient of increasing concentration of ammonium formate buffer (35 mM, 50 mM, 75 mM, 100 mM, 125 mM, 150 mM, 250 mM and 350 mM ammonium formate, 30% v/v ACN and 0.1% formic acid; pH = 2.9).  Peptides from each of these fractions were cleaned-up using C18 ZipTip (Millipore, USA). Each fraction was then analysed on a quadrupole-TOF hybrid mass spectrometer (TripleTOF 6600, Sciex, USA) coupled to a nano-LC system (Eksigent NanoLC-425). 4 µg of these peptides were loaded on a trap-column (ChromXP C18CL 5µm 120Å, Eksigent) and desalted at a flow rate of 10 µl per minute for 10 min. Peptides were then separated on a reverse phase C18 analytical column (ChromXP C18, 3µm 120 Å, Eksigent) using buffer A (98 % water with 0.1 % formic acid and 2 % acetonitrile) and buffer B (98 % acetonitrile with 0.1 % formic acid and 2 % water) and the following gradient: buffer B was increased gradually from 3% to 25% in the first 68 min. It was increased to 35% solvent B in the next 5 minutes, In the next 2-minute buffer B was ramped up to 80% and held at the same concentration for next 3 minutes. Buffer B was swiftly brought to an initial 3% concentration in the next 1 min and held at the same concentration till the end of 87 minutes gradient.

The optimized source parameters were as follows: the ion spray voltage was to 5.5 KV, curtain gas was set at 25 psi, nebulizer gas was set at 10 psi and source temperature was set at 200°C. For DDA, a 1.8 s instrument cycle was repeated in high sensitivity mode throughout the entire gradient, consisting of a full scan MS spectrum (400–1250 *m/z*) with an accumulated time of 0.25s, followed by 30 MS/MS experiments (100–1500 *m/z*) with 0.050 s accumulation time each, on MS precursors with charge state 2+ to 5+ exceeding a 150 cps threshold. Rolling collision energy was used and the former target ions were excluded for 15 seconds.

For the DIA-SWATH run, instrument and chromatographic conditions were identical to the DDA run for library preparation. 100 precursor isolation windows were defined based on precursor *m/z* frequencies in the DDA runs, with a minimum window width of 5 *m/z* using the SWATH Variable Window Calculator (Sciex). The accumulation time was set to 0.25 sec for the MS scan (400–1250 *m/z*) and 0.025 s for the MS/MS scans (100–1500 *m/z*) respectively. Rolling collision energies were applied for each window based on the *m/z* range of each SWATH and a charge 2+ ion, with a collision energy spread of five. The total cycle time was 2.84 s.

**Database searches and peak extraction**

The spectral ion library, a merge search for 8 DDA runs and analysis in protein pilot software v5.0.1 (Sciex, USA) with the paragon algorithm, to obtain protein identities. The parameters were set as follows: cysteine alkylation—IAA, digestion—trypsin. The search effort was set to ‘thorough ID’ and false discovery rate (FDR) analysis was enabled. Proteins identified with a 5% FDR were considered. The search was carried out against the UniProt database containing 20,350 human proteins. The result (.group) file, thus generated served as the spectral ion library.

SWATH generated data processing was performed using the SWATH Acquisition Microapp 2.0.1 in PeakView 2.1 Software. Protein Pilot search result file (.group) was imported with 251 specified proteins, and shared peptides were excluded. Retention time alignment was performed using peptides from abundant proteins. The processing settings for peak extraction were set as: 10 peptides per protein, 5 transitions per peptide, >95% peptide confidence threshold, 5%FDR and exclude modified peptides. The XIC extraction window was set to 85 min with a 50 ppm XIC width. All information was exported in the form of Marker View (.mrkw) file as described ( Basak et al., 2015 ) ^12^.
